# Fundamental constraints on the evolution of vertebrate life histories

**DOI:** 10.1101/2024.01.23.576873

**Authors:** George C. Brooks, Josef C. Uyeda, Nicholas Bone, Hailey M. Conrad, Christopher G. Mull, Holly K. Kindsvater

## Abstract

Vertebrate life histories evolve in response to selection imposed by abiotic and biotic environmental conditions while being limited by genetic, developmental, physiological, demographic, and phylogenetic processes that constrain adaptation. Despite the well-recognized shifts in selective pressures accompanying transitions among environments, identifying the conditions driving innovation and the consequences for life-history evolution remain an outstanding question. Here, we compare the traits of aquatic and terrestrial vertebrates to infer shifts in demographic and evolutionary constraints that explain differences in life-history optimization. Specifically, our results emphasize the reduced potential for life-history diversification on land, especially that of reproductive strategies. Moreover, our study reveals differences between the evolution of viviparity in the two realms. Transitions from egg laying to live birth represents a major shift across life-history space for aquatic organisms, whereas terrestrial egg-laying organisms evolve live birth without drastic changes in life-history strategy. Whilst trade-offs in the allocation of resources place fundamental constraints on the way life histories can vary, ecological setting influences the position of species within the viable phenotypic space available for adaptive evolution.

## INTRODUCTION

The evolution of diversity of vertebrate life histories has been fuelled by key transitions among water and land, and the evolution of innovative traits or reproductive strategies that allow species to survive in novel environments. The consequences of such innovations can have cascading effects on available niche space (i.e., arising from physiological constraints), life histories, and macro-evolutionary patterns. For example, internal fertilization is a necessary but insufficient innovation to explain both the colonization of terrestrial environments, and the repeated transitions to bearing live young (viviparity) [1,2,3,4]. Viviparity is intricately connected to parental care, egg size, and fertilization mode [4,5,6,7]. These characteristics, in turn, can limit evolutionary lability or allow species to capitalize on novel ecological opportunities [8,9,10,11]. The evolution of viviparity, for example, enables vertebrates to overcome some abiotic constraints, such fact that eggs without shells will desiccate on land, but will require increased investment in each offspring, shifting the fundamental trade-off between offspring size and number. These processes can affect the parameter space of life history traits that are viable, which underly demographic constraints, ultimately affecting the course of evolution. We can infer that the trait-space occupied by species persisting over evolutionary timescales emerges from these combined processes. We use this perspective to understand how transitions in reproductive mode have shaped the evolution of vertebrate diversity. While links between live-birth and adaptive radiation have been documented in fish [8], mammals [12], and reptiles [13], the consequences of the evolution of viviparity for life-history traits and demography in aquatic and terrestrial environments are not fully understood.

Life history traits related to growth, reproduction, and survival determine an individual’s maximum size, fitness, and pace-of-life and provide a fundamental link between population demography and growth as well as evolutionary dynamics [14,15,16,17]. Life-history diversity evolves according to the biotic stressors (e.g., predation, competition), characteristics of the environment, and the genetic and developmental context of each lineage [14,18,19]. The selection pressures of each environment dictate which life-history strategies persist over evolutionary time and render certain strategies impossible (e.g., extremely large body mass on land, broadcast spawning on land) [20,21], while phylogenetic inertia contributes by determining which strategies can be attained via evolution. Transitions between aquatic and terrestrial habitats have led to radiations into unexploited niches in both habitats. These transitions require key adaptations in development and reproduction that foster success in the new context. Alongside the preeminent colonization of land that marks the origin of tetrapods, transitions to an entirely terrestrial life history have evolved independently in multiple amphibian clades, accompanied by the evolution of direct development, in which juveniles complete metamorphosis prior to hatching on land [7]. Reversions back to the water have occurred in most vertebrate clades [22].

The fundamental differences between species in water versus on land reflect the consequences of the unique physical and ecological features of life in water for growth, survival, and reproduction [23,24,25,26]. Aquatic environments prevent desiccation, buffer thermal ranges relative to the extreme temperatures possible on land and offer structural support against the pull of gravity [23,24,25]. Additionally, aquatic food webs differ in their complexity and the spatial distribution of resources [27,28,29]. External fertilization is common among aquatic species, most of which release small eggs that disperse in the water column [23,25]. However, the repeated evolution of internal fertilization has made the transition to viviparity possible in some ancestrally egg-laying lineages of elasmobranchs (sharks and rays), teleosts (ray finned fishes), and amphibians [30]. A key innovation in vertebrate diversification on land was the evolution of amniotic egg, which requires internal fertilization, and prevents desiccation of eggs in air. Several amniotic vertebrates have subsequently evolved partially or fully aquatic life histories but must lay and care for their eggs on land. A few amniotic lineages, such as some sea snakes and aquatic mammals, have escaped this constraint by evolving viviparity [30]. While the sequence of parity mode evolution remains a topic of debate in squamate reptiles [30,31], this uncertainty mostly concerns the choice of theoretical and statistical model, and a more comprehensive framework with multiple lines of evidence from physiology, morphology, and developmental biology supports the general sequence of egg-laying to live-bearing with potential for reversals [32]. Egg-laying amniotes, such as some reptiles and all birds, can approximate viviparity by way of producing large eggs and providing extensive parental care, [7,33], but are prohibited from a fully aquatic existence.

Despite the well-recognized shifts in selective pressures accompanying transitions among environments, the conditions driving developmental innovation and its consequences for life history diversity among vertebrates have never been comprehensively tested. One challenge is the extensive macroevolutionary scale that is necessary to identify the constraints imposed by transitions among habitats or reproductive mode. At this scale, the traits that taxa use to modulate their life history may not be comparable across clades (e.g., parity mode may be a good proxy for parental investment in many squamates, but not in amphibians). This means that finding traits that allow quantitative comparisons of across clades can be challenging [34,35]. Here, we compared extant vertebrates, including cartilaginous fishes, bony fishes, amphibians, reptiles, birds, and mammals to demonstrate that the requirements of life on land, or the ecological release experienced when colonizing a novel environment, exert a predictable shift in demographic and evolutionary constraints, with important consequences for the evolution of biodiversity. To accomplish this, we define a novel trait space that enables comparisons of the complex, interrelated sets of trade-offs underlying all life histories [35,36]. Our use of broadly available trait data facilitates the generation of hypotheses and tests of these hypotheses with macroevolutionary data. Specifically, we use life-history traits that serve as proxies for mortality of juveniles and adults [37,38,39] to model the diversification of vertebrates, including during the vertebrate invasion of land, reversions back to the water, and after the evolution of live birth.

We identify a three-way trade-off on the bounds of the relationship between juvenile mortality, adult mortality, and body size (Supplementary Fig. 1-3). Using this trade-off, we propose an alternative phylogenetic framework for studying constraints on life history evolution in vertebrates. Given the 450-million-year history of vertebrate evolution, we assume that enough time has passed for the species in our dataset to fully explore the bounds of life-history space, i.e., areas of trait space where species are absent represent non-viable life-history strategies as opposed to trait space that species have yet to evolve into [40,41]. We reformulate the bounded state space of life history as a “Ternary plot” – equilateral triangles that define a space based on trade-offs in investment among these interacting variables [34,42]. Focusing on traits that are fundamental to fitness and placing each species in subdivided locations of this common space allows us to explore transitions among locations and quantify which direction of transitions are more common over evolutionary time. With this approach, we can go beyond previous research grouping species by traits in ternary space to formally quantify whether and how the evolutionary consequences of transitions from oviparity to viviparity differ among aquatic and terrestrial environments.

## RESULTS

### Life-history patterns in aquatic and terrestrial environments

We used phylogenetic comparative tests to identify constraints on the value of traits in life-history space. By focusing on defining these constraints, as opposed to variation in the rates of change, we uncovered fundamental shifts in life history across vertebrates due to transitions between aquatic and terrestrial life histories. Larger vertebrates have lower mortality, as expected, but for aquatic and terrestrial organisms this is achieved through different means (Fig. 1). Terrestrial vertebrates have low juvenile mortality regardless of body size but exhibit a steep negative relationship between body size and adult mortality (*β* = −0.04, *β* = −0.58 respectively, *p* < 0.001, Fig. 1). In contrast, species with aquatic offspring show a large reduction in juvenile mortality with increasing body size, and low adult mortality across taxa with a weak negative relationship with body size (*β* = −0.40, *β* = −0.17 respectively, *p* < 0.001, Fig. 1). When taken together, aquatic and terrestrial species exhibit highly concordant patterns of lifetime mortality with body size (*β* = −0.54, *β* = −0.57 respectively, *p* = 0.29, Fig. 1). Body size and environment explain over half of the variation (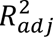 = 0.57) in juvenile mortality of vertebrates, and over one third of the variation in adult (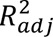 = 0.41) and lifetime mortality (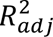 = 0.37).

**Figure 1:**
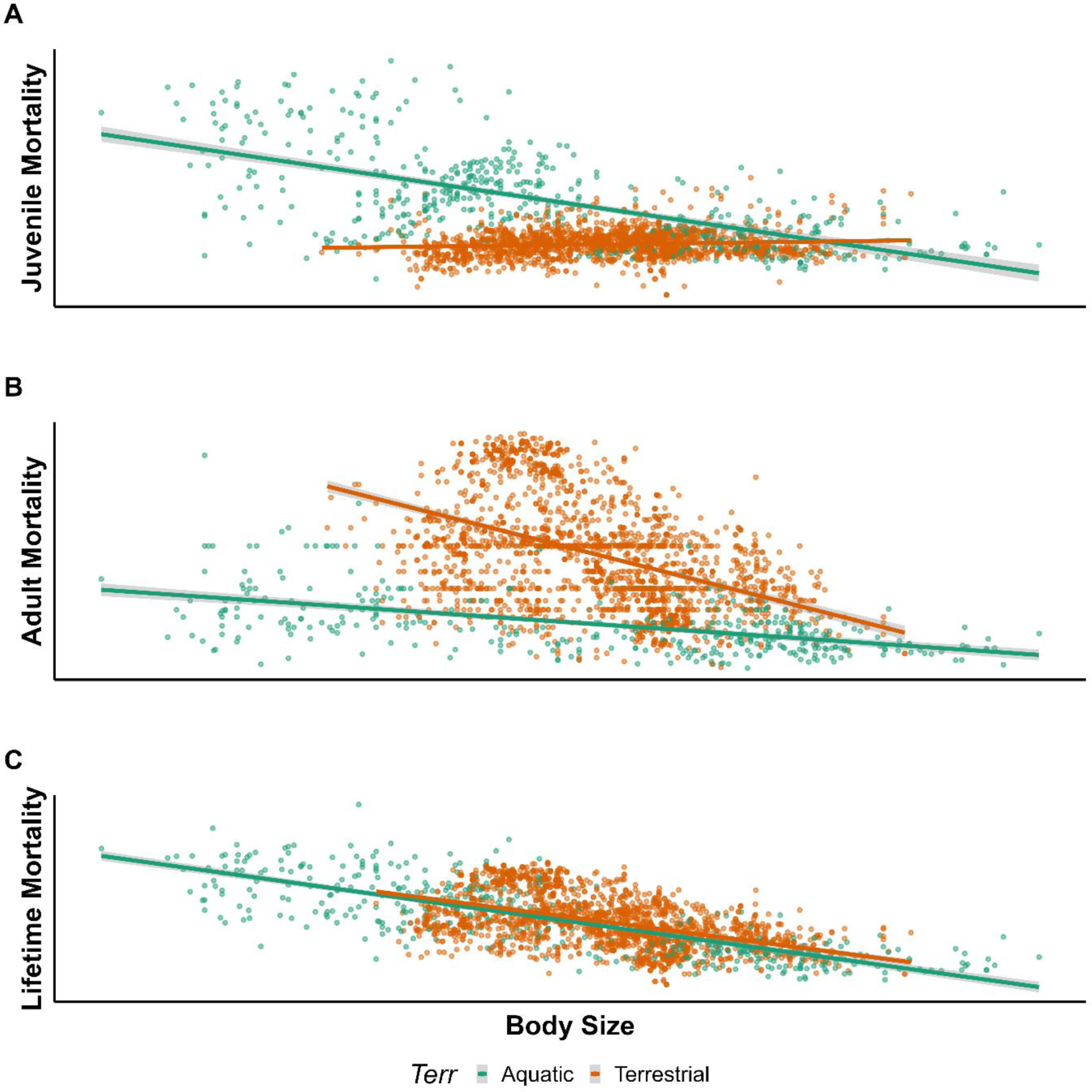
The relationship between body size and a) juvenile mortality, b) adult mortality, and c) lifetime mortality for aquatic (n = 577) and terrestrial (n = 2345) vertebrates. Aquatic and terrestrial species differ in their relationship between juvenile mortality and body size (*GLM*, *p* < 0.001), and in their relationship between adult mortality and body size (*GLM*, *p* < 0.001). When taken together however, aquatic and terrestrial species show no difference in the scaling of lifetime mortality with body size (*GLM*, *p* = 0.29). Lines represent predicted trends, and shaded areas reflect 95% confidence intervals.

These analyses additionally highlighted that key trade-offs among life history traits depend on the scale of comparisons [32,43,44]. For example, we found that non-phylogenetic estimates of the covariance between body size, adult mortality, and juvenile mortality are all strongly negative (Supplementary Table 1). While under most scenarios phylogenetic estimates are also negative, these estimates are considerably weaker than the non-phylogenetic ones (except size and adult mortality, which was roughly of similar in magnitude in the phylogenetic estimates, Supplementary Table 2). Furthermore, after accounting for the interaction between size and terrestrial habit, we found size and juvenile mortality to have positive evolutionary correlations in aquatic taxa, despite having strong negative correlations in the non-phylogenetic analyses (Supplementary Table 1). Differences between the non-phylogenetic and phylogenetic covariances suggest a contrast between the trade-offs operating in the evolutionary dynamics between closely related species (i.e. the covariance of changes in among diverging lineages) in life history traits with high phylogenetic signal (Supplementary Table 3) and the overall bounds on the relationship (the overall covariance among variables across species). This is apparent by visual inspection of the relationships between variables (Fig. 1 and Supplementary Fig. 3-5). This difference emerges because while evolutionary changes may appear positively or negatively correlated in the short term, the full space of life history variation in vertebrates is bounded by fundamental trade-offs that become visible in the non-phylogenetic regressions (and in examination of the life history space ordinated across species).

### Parity mode evolution in aquatic and terrestrial environments

When modelling life-history evolution in ternary space, we found differences in the evolutionary consequences of transitions to viviparity in each environment. Overall, aquatic species can occupy virtually every corner of life-history space, whereas terrestrial species have strong selection pressure to evolve toward reduced adult and juvenile mortality and larger body size (Fig. 2). In both environments, while oviparity does not appear to exert strong constraints on life-history strategies, transitions to viviparity tend to be associated with reduced adult and juvenile mortality and larger body sizes (Fig. 3). This shift is explained by movement to a common location within the life-history triangle, despite different starting points within oviparous ancestors, and is generally driven by the shifts to viviparity in aquatic species including cartilaginous fishes (sharks and rays), bony fishes, and amphibians (Fig. 3, 4, and 6). The strength of this pattern varies among these clades; for example, transitions to live-bearing in sharks and rays are associated with a larger increase in body size than in bony fishes [45]. For terrestrial species, viviparity is associated with a much smaller shift in life-history space (Fig. 5, 6). In reptiles, which have the greatest number of origins of viviparity in terrestrial vertebrates, life history traits vary only minimally between oviparous and viviparous species.

**Figure 2:**
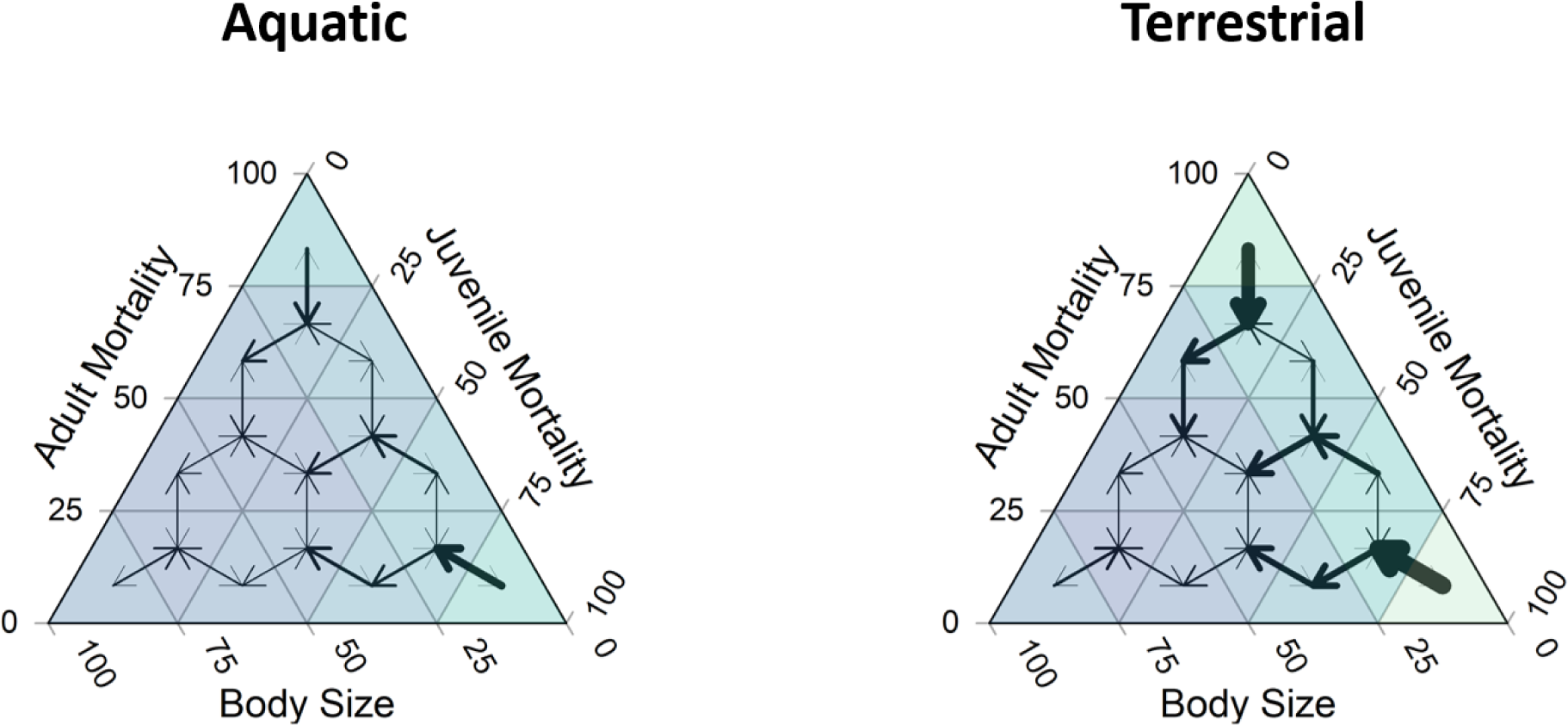
Movement through life-history space for aquatic and terrestrial vertebrates. Plots were generated from 2922 species for which complete life-history data was available. The colours represent the number of taxa represented in each segment, with darker colours representing more species, and lighter colours representing fewer species. The arrows indicate the direction and relative strength of selection through life-history space.

**Figure 3:**
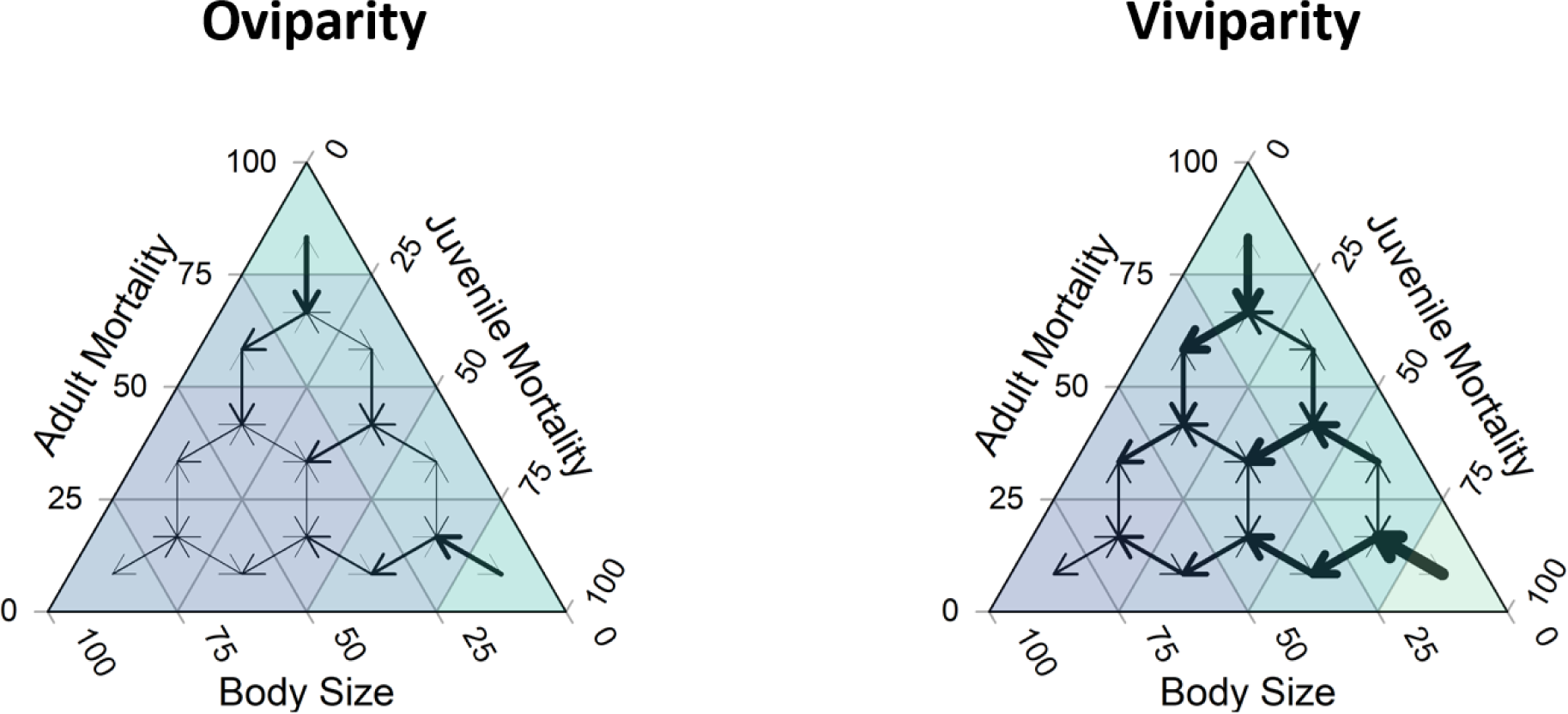
Movement through life-history space for egg-laying and live-bearing vertebrates. Plots were generated from 2922 species for which complete life-history data was available. The colours represent the number of taxa represented in each segment, with darker colours representing more species, and lighter colours representing fewer species. The arrows indicate the direction and relative strength of selection through life-history space.

**Figure 4.**
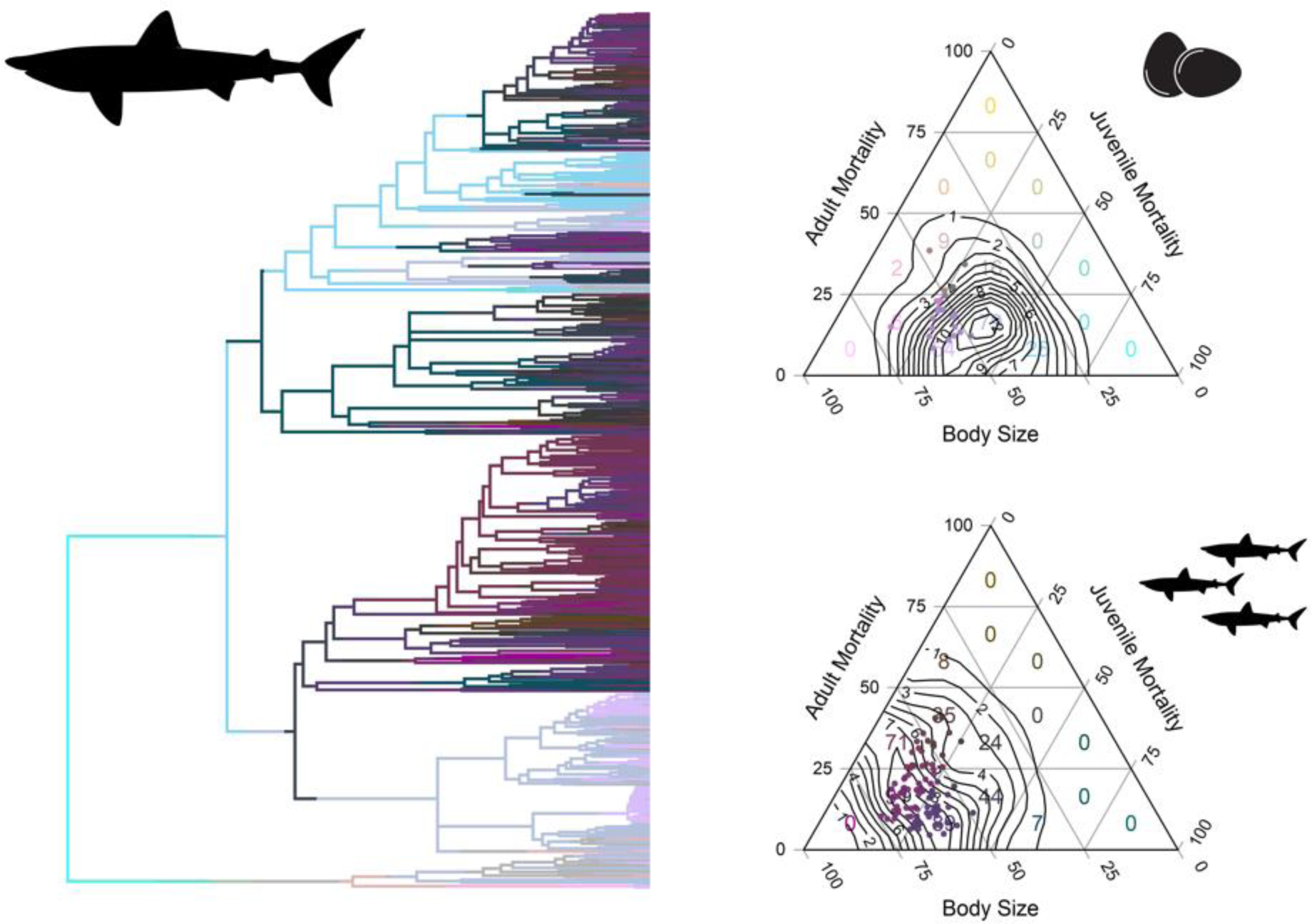
Phylogeny of cartilaginous fishes (sharks, rays, and chimaeras) and the relative position in life-history space for egg-laying (top right) and live-bearing (bottom right) taxa. Ternary plots show the counts of species (n = 561) in each segment with overlaid contours indicating the density of species in life-history space, with species with complete data indicated as points. Densities were estimated using the two-dimensional kernel density estimation procedure from the *Ternary* package [81]. The branches of the phylogeny are coloured using stochastic mapping to match the cells in the ternary plots, with lighter and darker colors indicating oviparous and viviparous species, respectively. Silhouettes were obtained from PhyloPic (https://www.phylopic.org/) under a creative commons license.

**Figure 5.**
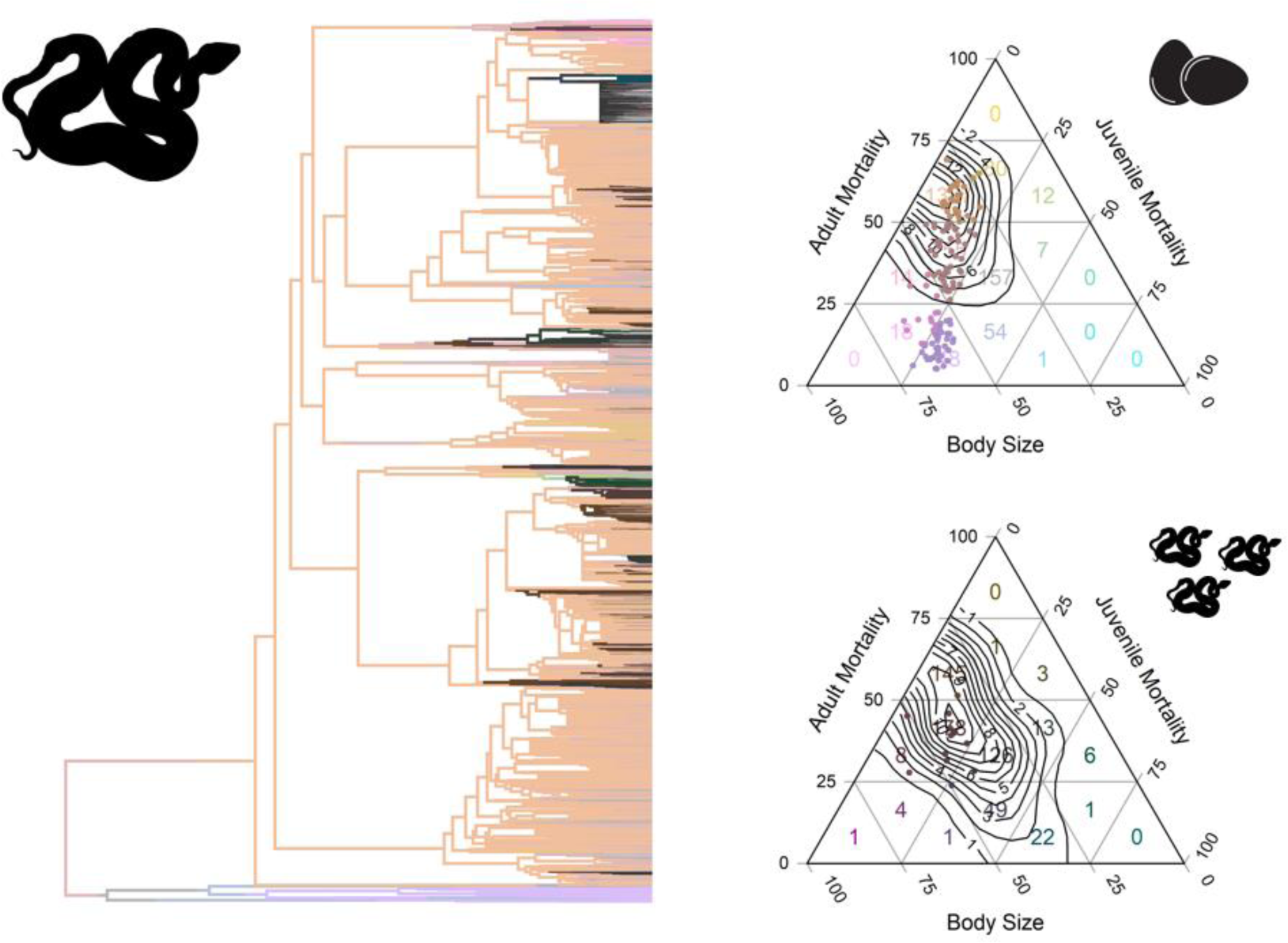
Phylogeny of reptiles and the relative position in life-history space for egg-laying (top right) and live-bearing (bottom right) taxa. Ternary plots show the counts of species (n = 3031) in each segment with overlaid contours indicating the density of species in life-history space, with species with complete data indicated as points. Densities were estimated using the two-dimensional kernel density estimation procedure from the *Ternary* package [81]. The branches of the phylogeny are coloured using stochastic mapping to match the cells of the ternary plots, with lighter and darker colors indicating oviparous and viviparous species, respectively. Silhouettes were obtained from PhyloPic (https://www.phylopic.org/) under a creative commons license.

**Figure 6.**
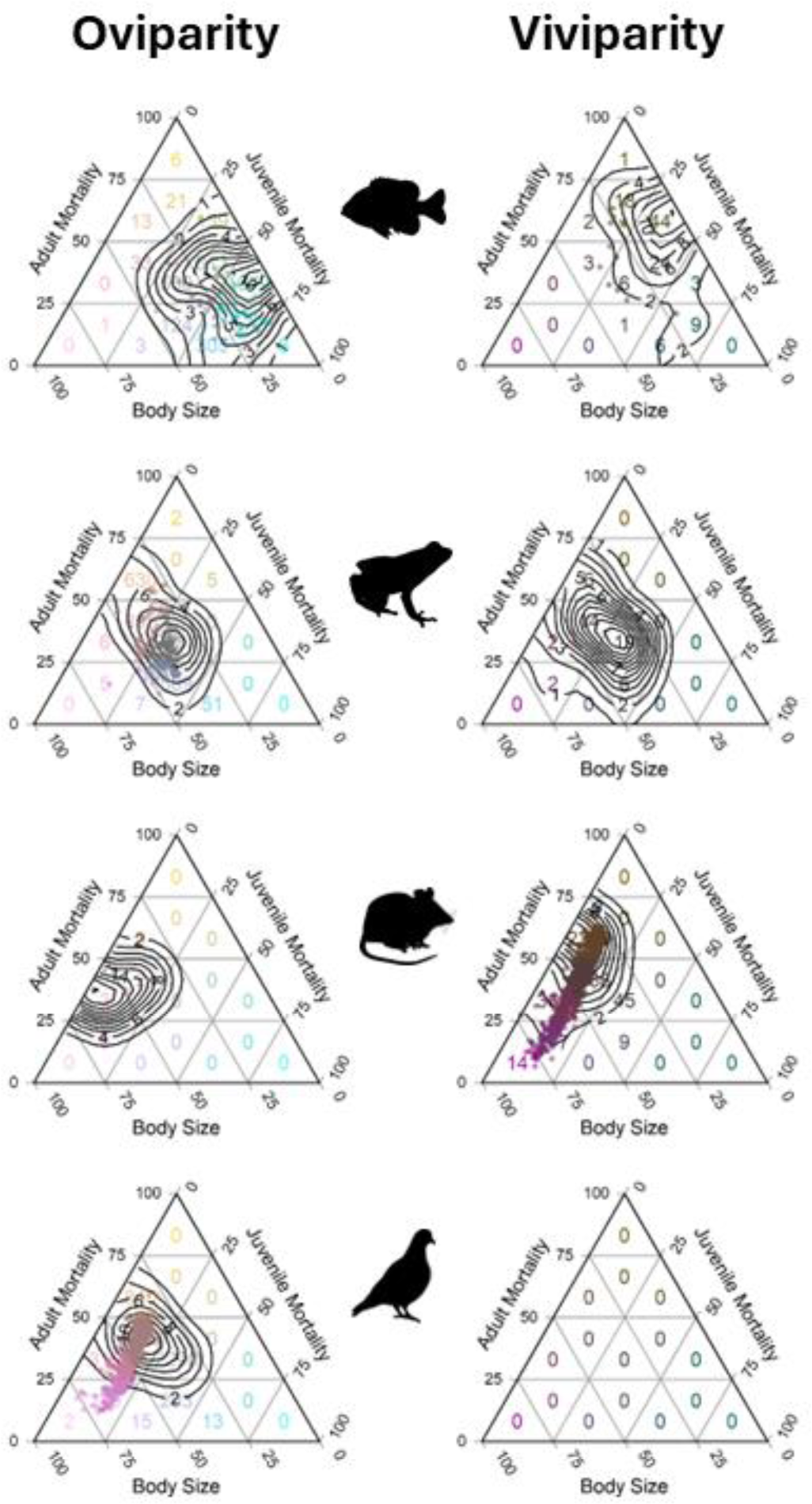
Comparison of egg-laying and live-bearing species of bony fish (n = 3799), amphibians (n = 4520), mammals (n = 4480), and birds (n = 8690). Ternary plots show the counts of species in each segment with overlaid contours indicating the density of species in life-history space, with species with complete data indicated as points. Densities were estimated using the two-dimensional kernel density estimation procedure from the *Ternary* package [81]. The branches of the phylogeny are coloured using stochastic mapping to match the cells in the ternary plots, with lighter and darker colors indicating oviparous and viviparous species, respectively. Silhouettes were obtained from PhyloPic (https://www.phylopic.org/) under a creative commons license.

## DISCUSSION

Here we present a novel framework for studying life history evolution that quantifies how the reproductive strategies of vertebrates have been shaped by contrasting fitness landscapes of aquatic and terrestrial environments. Specifically, transitions from egg laying to live birth have led to major shifts in life-history trait-space for aquatic organisms. By contrast, the life histories of oviparous and viviparous species in terrestrial environments are largely similar. Our analysis is based on a novel trait-based approach to classifying species based on three components of fitness (body size, age of maturity, and lifetime fecundity) which can inform comparative studies that seek to explain life-history evolution across a broad range of taxa. Our approach builds on previous research placing species in trait space to identify trade-offs shaping life history strategies, most notably the Equilibrium-Periodic-Opportunistic (E-P-O) categories [34,46]. Our method differs from prior work in that it relies on lifetime fecundity instead of offspring size as a proxy for juvenile mortality, and we use maturation age as a proxy of adult mortality. In our framework, lifetime mortality (the summed mortality of juvenile and adults) shows a remarkably consistent relationship with body size (Fig. 1), as expected, justifying our use of these proxy traits. Elsewhere, our approach has been used to identify Periodic and Opportunistic life histories, and to divide Equilibrium species into small-bodied species that have relatively few, large offspring (Precocial species), and large-bodied species, which may produce low numbers of large offspring but also live a long time (Survivor species) [16,37]. This distinction captures an important range of life history variation in mammals, despite their low numbers of offspring when compared to amphibians and fishes. Our focal traits are also closely connected to a species’ placement on the slow-fast life history continuum [19,47]. Our approach requires less knowledge of age-specific vital rates to place species in ternary space. Ultimately, our three metrics simplify life-history space over macroevolutionary time and highlight long-term constraints on the evolution of vertebrate diversity.

Overall, the selection pressures that favour live birth, as well as the benefits conferred, are different in the water versus on land. In aquatic environments, the evolution of viviparity requires the precursory evolution of internal fertilization, which is a prerequisite of life on land [3,48,49,50]. The origins and advantages of different modes of fertilization are not fully known; female choice, male opportunity to manipulate female investment, sperm storage, increased control over fertilization timing, and reduced egg mortality have all been suggested drivers of the evolution of internal fertilization [51,52]. The idiosyncrasies of development and early life history of aquatic species that have evolved along with externally fertilized eggs are likely a barrier to the evolution of viviparity for most aquatic vertebrates. Aquatic vertebrates with external fertilization produce many tiny offspring, which is hypothesized to maximize fitness when prey encounter is stochastic and survival is independent of density [53,54]. By contrast, viviparity in aquatic species with internal fertilization acts to increase juvenile survival, presumably through reduced predation of offspring held *in utero*. Terrestrial vertebrates have internal fertilization and exhibit life-history strategies that effectively minimize juvenile mortality regardless of whether they have eggs or bear live young. For example, terrestrial egg deposition requires larger eggs to prevent desiccation. Larger eggs require a longer period to develop, which in turn favours extended parental care. Juvenile animals on land are not bathed in a nutrient broth as they are in aquatic systems, thus precluding the evolution of particularly small offspring. Evolving viviparity on land does not appreciably reduce juvenile mortality, but instead could act to maintain high levels of juvenile survival across a wider range of environmental conditions.

Differences in life-history diversity and evolution in water versus on land carry important implications for the adaptive capacity of species to environmental change [55,56]. Firstly, terrestrial environments constrain the diversification of life-history strategies. As a result, terrestrial vertebrates will be more limited than aquatic species in their capacity to adapt to global change. Secondly, when considering the relative difference in vulnerability to extinction between egg-laying and live-bearing taxa, environmental considerations are paramount. Terrestrial live bearers have similar life-history traits to their egg-laying counterparts, and thus will show comparable response to anthropogenic threats, all else being equal. In contrast, the large shift in life-history strategies seen as viviparity evolves from oviparity in aquatic species suggests their response to threats may be starkly different. For cartilaginous fish, the evolution of live birth represents a distinct shift to a ‘slower’ life history strategy, increasing generation time and reducing their adaptive capacity to rapid environmental change. For bony fish, however, live birth appears to have evolved --when possible-- to buffer juvenile stages from highly stochastic, ephemeral environments, and is often accompanied by small body size and ‘faster’ life histories. The adaptive capacity of aquatic livebearers with short generation times in the face of anthropogenic threats may be greater than oviparous species.

Strong linear relationships (Fig. 1) and a triangular distribution of life-history strategies (Supplementary Fig. 3) indicate that long term macroevolutionary patterns are highly directional and suggest that three axes are sufficient to define the vertebrate trait space. Phylogenetically corrected analyses showed less clearly defined trade-offs between the three axes (Supplementary Fig. 4-6), suggesting that evolution over the short term is not constrained by classic life-history trade-offs between growth, survival, and reproduction [44,57], exemplified by the ubiquitous positive relationship between body size and clutch size within clades [58]. Individual clades may have different constraints, but they are all limited to this common space. In other words, a ternary plot is naturally bounded and inherently accounts for the trade-offs in resource allocation that every living organism must make [36]. Indeed, by defining the common life-history space of all vertebrates we provide a useful way to summarize the limits of evolution within a clade. Although the overarching constraint is the inability for species to evolve into ‘Darwinian demons’ which do not face trade-offs between growth, reproduction, and survival, here we illustrate how life-history space is filled by different clades in diverse ways. Only by understanding the macroevolutionary trade-offs that define trait space does the challenge of comparing such a disparate array of organisms become tractable; disparities in how species move through space can be used to answer questions about which traits are key levers controlling diversification.

Fundamentally, placing species in a common life-history framework helps us to understand the rules of life. Once all species are mapped in trait space, one can model macroevolutionary dynamics and explore covariance between key life-history traits. For example, we show that the evolution of viviparity in aquatic and terrestrial environments proceeds through different paths and at different rates. As an initial first step, we tested models where transitions happened in these traits independently of their location in state space. However, an interesting alternative model is one where each location in life history space has its own transition rate. Future elaborations of our approach could treat the evolution of viviparity as a cause, effect, or interaction between other factors. Additionally, our approach could be used to understand how differences in food web structure in aquatic and terrestrial environments shapes life-history strategies. Productivity, the patchiness of resources, and the distribution of predators in space will have profound implications for both a species’ transitions in life-history space and the position of life-history optima. Lastly, a potentially fruitful application of this work is to species’ invasions, for which it is possible to predict shifts in trait space following the introduction of a non-native species. If species are typically on the edge of replacement, they will likely not be able invest in one of the trait axes (larger body size, higher survival, increased reproductive output) without some kind of ecological release that temporarily relaxes existing constraints. The colonization of land and the ability of viviparous species to exploit cold climates both exemplify the interplay between life-history evolution and ecological opportunity. These examples demonstrate how unifying classic life history theory with trait-based approaches and contemporary computational techniques can be leveraged to make progress on previously intractable questions and generate novel insights into the forces that speed up or slow evolutionary innovation.

## METHODS

### Trait-based framework

We collated published data on species’ body size, age at maturity, and lifetime fecundity, allowing us to place species on common life-history axes, despite differences in their ecology. We compared distribution of body sizes, age at maturity, and lifetime fecundity across vertebrate classes (Supplementary Fig. 1). We related this combination of traits to live-bearing (viviparous) and egg-laying (oviparous) reproductive strategies (Supplementary Fig. 2), and the occupancy of aquatic and terrestrial environments (Supplementary Fig. 3). Several vertebrate groups (e.g., amphibians, sea turtles) occupy multiple environments during their lifetimes, but for simplicity we classified species as aquatic or terrestrial based on the initial environment a species inhabits (egg deposition site for oviparous species, juvenile environment for viviparous species). Thus, most amphibians are classified as aquatic despite extended terrestrial phases and sea turtles are classified as terrestrial owing to their use of nesting beaches. The choice to focus on the environment of early-life stages stems from our interest in the evolution of reproductive strategies (e.g., parity mode) and the observation that fecundity exhibits the greatest variability and disparity across aquatic and terrestrial realms (Supplementary Fig. 1-3). Whilst a model that includes both juvenile and adult environments would be more informative, it is computationally infeasible to include all possible life cycle transitions in the state-space approach presented here.

Like other trait-based analyses of trade-offs and patterns in life histories [19,21,34], our trait proxies were derived from life-history theory [14,15]. However, our approach differs in that it uses only three traits to represent vital rates that are fundamental to both individual fitness and population demography, allowing us to maximize the number of taxa that we included in our analysis. Furthermore, unlike prior studies, we did not control for body size when comparing birth and death rates, instead addressing explicitly its changes after evolutionary transitions in habitat or reproductive mode. We relied on the principle that persistence of populations and species over evolutionary timescales must all fulfil the same requirement, namely that individual females, on average, must replace themselves and their mate [18,35,36]. Lifetime production of offspring is expected to be positively correlated with juvenile mortality and negatively correlated with per-offspring parental investment [35,37]. Similarly, age at maturity is strongly linked to adult mortality, whereby species that mature later have higher survival and longer expected lifespans [38,39]. Organisms with longer lifespans therefore must, on average, replace themselves at a slower rate than those with shorter lifespans; any other scenario would result in the unrealistic outcomes of exponential population growth or extinction. Both adult and juvenile mortality are also strongly related to body size, with larger animals often exhibiting larger clutch sizes and delayed maturity [40,41]. Fundamentally, body size represents investment that must be made in the growth and metabolic maintenance of body tissues outside of direct investment in reproduction.

### Trait and phylogenetic data

We assembled a phylogeny for vertebrates on an order-level backbone obtained using median divergence times from the timetree of life [59]. A supertree was made by grafting published trees for each clade. We combined trees for fish [60], mammals [61,62], birds [63], squamate reptiles [64], amphibians [65], turtles (testudines) [66], crocodilians [67], and chondrichthyans [68]. As the full chondrichthyan phylogeny was constructed with a birth-death polytomy resolver, we chose to base our analyses on the molecular backbone phylogeny calibrated with 10 fossil calibration dates comprising 610 species from all major orders. The phylogeny includes both oviparous and viviparous lineages and provides good coverage for the order Carcharhiniformes, which is the largest order of sharks and exhibits the greatest diversity of reproductive modes in this group.

Life history traits were compiled from publicly available databases that cover the major vertebrate clades (Table 1). A link to the complete dataset for all vertebrate clades used in this paper is provided in the data availability statement, and graphically displayed in the supplementary material (Supplementary Fig. 7). Data for bony fish (Actinopterygii) were downloaded from FishBase using the rfishbase package (www.fishbase.org) [69] and data for viviparous species was supplemented with the primary literature. Data for sharks and rays (Chondrichthyans) were taken from the Sharkipedia data repository (www.sharkipedia.org) [70]. Amphibian trait data were taken from AmphiBio [71]. Data for squamate reptiles were taken from [72] and supplemented with estimates of longevity sourced from the literature. Data for birds and mammals were obtained from [73]. Manual searches of the literature were conducted and bolstered our dataset for an additional 17 species of bony fish, 137 snakes, 137 amphibians, 45 turtles and 9 crocodilians. Owing to the number of species included in the analysis, we had to assume that the information obtained from previously published datasets were reliable but retrospectively checked outliers from visual inspection of the model output to confirm their accuracy.

**Table 1.** Number of species for each of the major vertebrate clades for which partial and complete life-history data was available. Numbers in parentheses show the proportion of total diversity represented within each clade. Also shown is the proportion of species (with complete data) that are viviparous or have terrestrial egg deposition.

| **Clade** | **Partial Data** | **Complete Data** | **Viviparity** | **Terrestrial Eggs** |
| --- | --- | --- | --- | --- |
| **Chondrichthyes** | 571 (46%) | 115 (19%) | 81% | 0% |
| **Actinopterygii** | 9441 (29%) | 117 (0.3%) | 15% | 0% |
| **Amphibia** | 5250 (61%) | 260 (4%) | 3% | 6% |
| **Reptilia** | 4516 (39%) | 213 (3%) | 14% | 100% |
| **Aves** | 8690 (80%) | 922 (8%) | 0% | 100% |
| **Mammalia** | 4480 (70%) | 1465 (30%) | 99% | 97% |

Body size measurements are reported as snout-vent length (SVL) for all amphibians, reptiles, and mammals. Body sizes reported for bony fish, sharks, skates, and chimaeras represent total length, and measurements for rays are disc (wing) width. When multiple measurements were available for a species, mean values were used for analysis. If sex-specific estimates were available, we used numbers reported for females. Most trait databases did not include mass, so to convert lengths to mass we used available female length-weight (L-W) or width-weight conversion relationships (see supporting data for sources of L-W relationships). When sex and species-specific L-W relationships were unavailable, we estimated reasonable L-W relationships from the same genus or family.

Reproductive mode was characterized as either egg-laying or live-bearing, and fertilization mode was coded as internal or external. Species were classified as either aquatic or terrestrial; given our interest in reproductive modes, species with biphasic life cycles were coded based on the location of egg deposition (e.g., most amphibians are coded as aquatic despite having terrestrial adults). Given our emphasis on the evolution of reproductive strategies, we only considered juvenile environments in the present analysis.

Proxies for mortality can be inferred from the life-history traits that are most sensitive to changes in mortality schedules [14]. For example, increased predation or harvest typically selects for earlier maturity [74,75]. Therefore, we use the inverse of age at maturity to estimate adult mortality, such that:

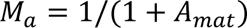

where *A*_*mat*_ is the age at maturity and *M*_*a*_ is adult mortality [37]. In a similar fashion, lifetime fecundity is indicative of mortality experienced in early life-stages [14,15]. Calculating lifetime fecundity requires information on lifespan, age at maturity, breeding frequency (e.g. annual, biennial) and clutch size. We logged this quantity such that lifetime egg production is equal to:

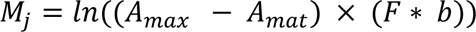

where *A*_*max*_is longevity, *A*_*mat*_is maturity, *F* is fecundity, and *b* represents the number of breeding bouts per year. The distribution of all three life-history traits, split between aquatic and terrestrial species, with egg-laying and live-bearing reproductive modes is presented in Supplementary Fig. 1 and summarized in Supplementary Table 4. To produce a rough measure of lifetime mortality for each species, we scaled and summed our proxies of juvenile mortality and adult mortality.

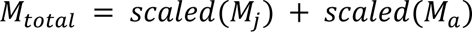

where our proxies of juvenile and adult mortality are first scaled before they are summed.

### Statistical Analyses

We estimated the covariances among traits using both phylogenetic and non-phylogenetic estimates using multivariate Brownian Motion models of trait evolution. The assumption of multivariate Brownian Motion is likely inappropriate, and therefore we also used an alternative phylogenetic modelling approach (Supplementary Table 2). Nevertheless, estimates of phylogenetic and non-phylogenetic correlation matrices among traits are a meaningful summary statistic of the covariances among traits across the observable evolutionary process. Therefore, we used a Pagel’s λ model and the R package *phylopars* [76] to accommodate for partially missing data, with analyses run multiple times with species missing 0, 1, or 2 trait values excluded. Analyses were performed once with a fully phylogenetic model (λ = 1) and again in a non-phylogenetic model (λ = 0) to facilitate comparisons of the phylogenetic and non-phylogenetic estimates with missing data. For each trait individually, we additionally report estimated phylogenetic signal metrics of Pagel’s λ and Blomberg’s K for all taxa with complete data using the function *phylosig* in the R package *phytools* [77].

To determine the relationship between maximum body size and mortality estimates, we fit three separate linear regression models with juvenile mortality, adult mortality, and lifetime mortality as response variables. In each model body size, habitat (aquatic or terrestrial), and their interaction were included as predictors. As we were interested in the overall bounds on life-history space, rather than clade-specific relationships, our primary analyses did not correct for phylogeny – the uncorrected formulation is likely more informative for our research question as it emphasizes the bounds of the trait space rather than the covariances of changes among lineages [32,43,44]. We then performed the same regressions accounting for phylogeny using the function *phylopars.lm* [76]. We assumed phylogenetically correlated residuals evolving under an Ornstein-Uhlenbeck process (Supplementary Fig. 5). For each model we obtained average effect sizes, and their associated standard errors and p-values. Model fit was evaluated by comparing predicted and observed mortality values and estimating *R*^2^ using the *rr2* package [78].

Ternary plots are often used for “phases” or regions of interactions that have a distinct shared feature (e.g. ‘solid’, ‘liquid’ or ‘gas’ equilibria in a Gibbs triangle). These features make them useful for projecting relationships between life history traits (e.g., Mims et al. 2010). We created ternary plots for our proxies for body size, adult mortality, and juvenile mortality. Allocation to each of these trait axes for each species was taken as their relative location across the full span of vertebrate life history trait space, with each of these ranges standardized to a new axis on a 0-100 scale [100 × (*x* − *x*_*min*_)⁄(*x*_*max*_ − *x*_*min*_)]. We then discretized this space into 16 equal triangles (4 bins for each trait) representing the possible states that each species in our dataset could occupy. This defined life-history space enables us to employ a discrete-state evolutionary model. By doubling the state space and allowing transitions between “layers” to be defined by binary characters, we can compare how such binary factors influence life history evolution across vertebrates. For example, we defined a 32-state continuous-time Markov chain with two ternary plot spaces corresponding to aquatic and terrestrial life history (Supplementary Fig. 8).

We then estimated transitions between aquatic and terrestrial environments, and between regions of the life history space assuming that movement within this space is only allowed between discrete bins that neighbor each other with a shared edge. Transitions in the binary character move species to corresponding bins in the replicated ternary space for the alternative binary state (Supplementary Fig. 8). Since a 32-by-32 transition matrix would have an exceptionally large number of parameters under most models, we simplified movement within each ternary space by defining four parameters governing the dynamics of trait change. These parameters are defined by the directional trends toward each of the three vertices of space, as well as an overall parameter governing whether species are attracted or repelled to the center of the triangle. Each layer of the life history space is defined by four unique parameters plus two parameters for asymmetric transitions in the binary trait, for a total of 10 parameters.

We conducted a second set of analyses that accommodate missing data in our three life history proxies by coding their location within the state space with ambiguous coding, which can be modeled using continuous-time Markov Chains. When modeled in this way, the posterior probabilities of species with missing data are computed jointly and informed by both their observed life history values, as well as the life history values of related species (see supplementary material). All analyses were performed in R using the *fit_mk* function in the castor package modified to account for ternary space [79](R Core Team 2023).

## Data availability

The dataset used in this study can be accessed at https://doi.org/10.5281/zenodo.11457349 [80]

## Code availability

The code generated in this study can be accessed at https://doi.org/10.5281/zenodo.11457349 [80]

## Acknowledgements

We would like to thank Jacob Berman and Ty Stephenson for help compiling trait data. We would like to thank the Department of Fish and Wildlife Conservation and the Department of Biological Sciences at Virginia Tech for financial and logistical support.

## Author contributions

GCB and HKK conceived the idea. GCB, HMC, and CM compiled and proofed the data. JU designed the analyses. GCB, JU, and NB performed the analyses and confirmed the reproducibility of the code and results. GCB, JU, and HKK wrote the manuscript. All authors revised and approved the final manuscript.

## Additional Information

Supplementary Information is available for this paper.

