## Supplemental Material for "Fundamental constraints on the evolution of vertebrate life histories"

### SUPPLEMENTARY TEXT

#### Supplementary methods

To fit a life history space to comparative data, we built a model in which our three life history variables are standardized by the range of observed values across vertebrates into a compositional simplex trait for adult mortality (*A*), juvenile mortality (*J*) and log body size (*M)* for species *i*, $S_i = \{A_i, J_i, M_i\}$ where $\sum S_i$ = 1. We then binned each dimension into *K* discrete bins, resulting $K^2$ discrete states, with the coordinates of the midpoint of each bin *k* given by the 3-state simplex $\boldsymbol{B^k} = \{B_A^k, B_J^k, B_M^k\}$. Transition rates between neighboring bins were calculated based on 4 directional rate parameters, $r = \{r_A, r_J, r_M, r_C\}$ and a rate scalar *R.* These rate parameters translate to transition rates from neighboring bins of index *i* to *j* of the instantaneous $K^2$ x $K^2$ transition matrix, **Q**, of the Markov process. If bins i and j are not neighbors in life history space, then $q_{ij} = q_{ji} = 0$ . However, if bins share an edge, then the transition rate, $q_{ij}$, is defined according to the equation:

$$q_{ij} = Re^{r_A(B_A^j - B_A^i) + r_J(B_J^j - B_J^i) + r_M(B_M^j - B_M^i) + r_C \Delta C} ,$$

where $\Delta C = \sum(\boldsymbol{B^j} - \boldsymbol{C})^2 - \sum(\boldsymbol{B^i} - \boldsymbol{C})^2$, or the difference in squared distance from the center of the ternary coordinate system, ***C* = *{1/3, 1/3, 1/3}*,** between bins *j* and *i*. In other words, *R* represents the overall rate scalar, $r_A$, $r_J$ and $r_M$, correspond to the coefficients for the bin center's change in each of the three simplex traits *A, J*, and *M* respectively, and $r_C$ represents the central tendency that either attracts or repulses evolution from the center of life history space**.** Note that there is some redundancy in this formulation of the parameters, and only two of the three values $\{r_A, r_J, r_M\}$ would be required to unique define a gradient of relative transitions values in **Q.** However, we obtained faster optimization and convergence to the maximum likelihood solution for values of **Q** in our preliminary trials by overparameterizing the model and including all 3 *r* values. These can then be algebraically reduced to only the 2 identifiable parameters.

For a binary trait predictor, *P*, such as oviparity vs. viviparity or aquatic vs. terrestrial, the total state space of the model is doubled to 2 x ($K^2$ x $K^2$) states. Each combination of traits *A, J*, & *M* can therefore be present in two bins, depending on whether *P = 0* or *P =1.* We index these bins as $\boldsymbol{B}_P^k = \{B_A^k, B_J^k, B_M^k\}$, where transitions can occur between corresponding bins $\boldsymbol{B}_0^k \rightarrow \boldsymbol{B}_1^k$ and $\boldsymbol{B}_1^k \rightarrow \boldsymbol{B}_0^k$ according to estimated transitions rates $q_{p,0\rightarrow 1}$and$q_{p,1\rightarrow 0}$, respectively. Each "layer" of life history space corresponding to either *P = 0 and P = 1* has freely estimated independent rate parameters, $R^P$ and $r^P$, doubling the number of rate parameters from 5 to 10, plus the two transition rates between layers, resulting in a total of 12 parameters in the model.

Simultaneous changes in $k$ and $P$ are disallowed meaning that all transition rates from $\boldsymbol{B}_0^k \rightarrow \boldsymbol{B}_1^{n \neq k}$ and $\boldsymbol{B}_1^k \rightarrow \boldsymbol{B}_0^{n \neq k}$ are set to 0.

### SUPPLEMENTARY TABLES

#### Supplementary Table 1. Correlation matrices estimated using *Rphylopars.*

| Max number of missing traits per species | Sample size | Correlation (95% CI) Size:AdMort | Size:JuvMort | AdMort:JuvMort |
|---|---|---|---|---|
| *Non-phylogenetic* (λ=0) | | | | |
| 0 | 2,921 | -0.31 (-0.37, -0.26) | -0.37 (-0.43, -0.31) | -0.17 (-0.21, -0.12) |
| 1 | 4,449 | -0.15 (-0.19, -0.12) | -0.55 (-0.63,-0.49) | -0.24 (-0.29, -0.20) |
| 2 | 25,071 | -0.12 (-0.16,-0.09) | -0.60 (-0.66, -0.55) | -0.24 (-0.29, -0.20) |
| *Brownian Motion* (λ=1) | | | | |
| 0 | 2,921 | -0.10 (-0.15, -0.064) | -0.042 (-0.082, -0.0052) | -0.068 (-0.11, -0.030) |
| 1 | 4,449 | -0.13 (-0.16, -0.094) | -0.043 (-0.084, -0.0053) | -0.079 (-0.12, -0.042) |
| 2 | 25,071 | -0.27 (-0.35, -0.20) | -0.081 (-0.18, 0.0086) | -0.059 (-0.10, -0. 016) |

**Supplementary Table 2. Phylogenetic regressions of adult and juvenile mortality against size x terrestriality using an Ornstein-Uhlenbeck model for the phylogenetic residual variation.**

| Max number of missing traits per species | Sample size | Response | Regression coefficients for scaled variables^ | | | |
|---|---|---|---|---|---|---|
| | | | Aquatic Intercept | Terrestrial effect† | Aquatic $\beta_{size}$† | Terrestrial $\beta_{size}$‡ |
| 0 | 2,921 | Juvenile Mortality | 0.212 | 0.255*** | 0.225** | -0.197*** |
| 1 | 4,449 | | 0.240 | 0.229*** | 0.195*** | -0.199*** |
| 2 | 25,071 | | 0.345 | 0.0496*** | 0.00222 | -0.100*** |
| 0 | 2,921 | Adult Mortality | 0.600 | -0.0163 | -0.597*** | -0.466 |
| 1 | 4,449 | | 0.636 | 0.148 | -0.637*** | -0.569 |
| 2 | 25,071 | | 0.428 | 0.169*** | -0.213*** | -0.499*** |

^All three variables scaled between 0 and 1 based on the range of minimum and maximum values in the dataset as described in the main text methods.
†Null hypothesis: Effect = 0; P-value < (*0.05; ** 0.01; ***0.001).
‡Null hypothesis: Coefficient significantly different from Aquatic $\beta_{size}$ (i.e. significant interaction).

**Supplementary Table 3. Phylogenetic signal estimates for life history variables. Significance was assessed from 100 randomization replicates using *phylosig* in the R package *phytools. Phylogenetic signal metrics reported are Pagel's λ and Blomberg's K.***

| Trait | λ | K | p-value | N |
|---|---|---|---|---|
| Body Size | 0.999 | 0.687 | $p < 0.05$ | 4,267 |
| Adult Mortality | 0.988 | 0.291 | $p < 0.05$ | 4,449 |
| Juvenile Mortality | 0.989 | 0.197 | $p < 0.05$ | 3,103 |

**Supplementary Table 4. Summarized life-history traits for each vertebrate clade.** Statistics are separated by parity mode and whether eggs are terrestrial or aquatic. Numbers reported are medians with the range in parentheses

| Trait | Class | Reproductive Mode | | | |
|---|---|---|---|---|---|
| | | **oviparous + aquatic** | **viviparous + aquatic** | **oviparous + terrestrial** | **viviparous + terrestrial** |
| **Body Size (cm)** | *Chondrichthyes* | 80.0 (32.0-253) | 110.0 (20.5-2100) | - | - |
| | *Actinopterygii* | 11.5 (0.8-360) | 6.5 (1.6-82.4) | - | - |
| | *Amphibia* | 4.5 (1.1-136) | - | 3.9 (1.1-130) | 21.7 (1.9-80) |
| | *Reptilia* | - | - | 6.2 (1.9-800) | 7.9 (3.5-600) |
| | *Aves* | - | - | 35.5 (9.3-158.8) | - |
| | *Mammalia* | - | 362.9 (34.9-3049) | 54.7 (37.5-67.5) | 18.5 (3.1-750) |
| **Juvenile Mortality** | *Chondrichthyes* | 5.3 (2.3-10.0) | 4.3 (2.3-8.2) | - | - |
| | *Actinopterygii* | 13.9 (2.9-21.0) | 4.6 (2.5-16.4) | - | - |
| | *Amphibia* | 8.1 (0.7-12.6) | - | 4.8 (2.9-8.1) | 4.1 (2.6-6.5) |
| | *Reptilia* | - | - | 4.5 (0.1-9.5) | 4.2 (0.1-6.7) |
| | *Aves* | - | - | 4.0 (0.6-7.2) | - |
| | *Mammalia* | - | 2.6 (1.5-4.0) | 3.5 (3.3-3.7) | 3.3 (0.1-6.5) |
| **Adult Mortality** | *Chondrichthyes* | 0.11 (0.06-0.31) | 0.13 (0.01-0.5) | - | - |
| | *Actinopterygii* | 0.25 (0.04-0.99) | 0.38 (0.04-0.92) | - | - |
| | *Amphibia* | 0.33 (0.09-0.83) | - | 0.25 (0.07-0.80) | 0.25 (0.14-0.57) |
| | *Reptilia* | - | - | 0.43 (0.02-0.91) | 0.30 (0.09-0.75) |
| | *Aves* | - | - | 0.50 (0.01-0.86) | - |
| | *Mammalia* | - | 0.12 (0.05-0.73) | 0.38 (0.36-0.40) | 0.53 (0.07-0.94) |

### SUPPLEMENTARY FIGURES

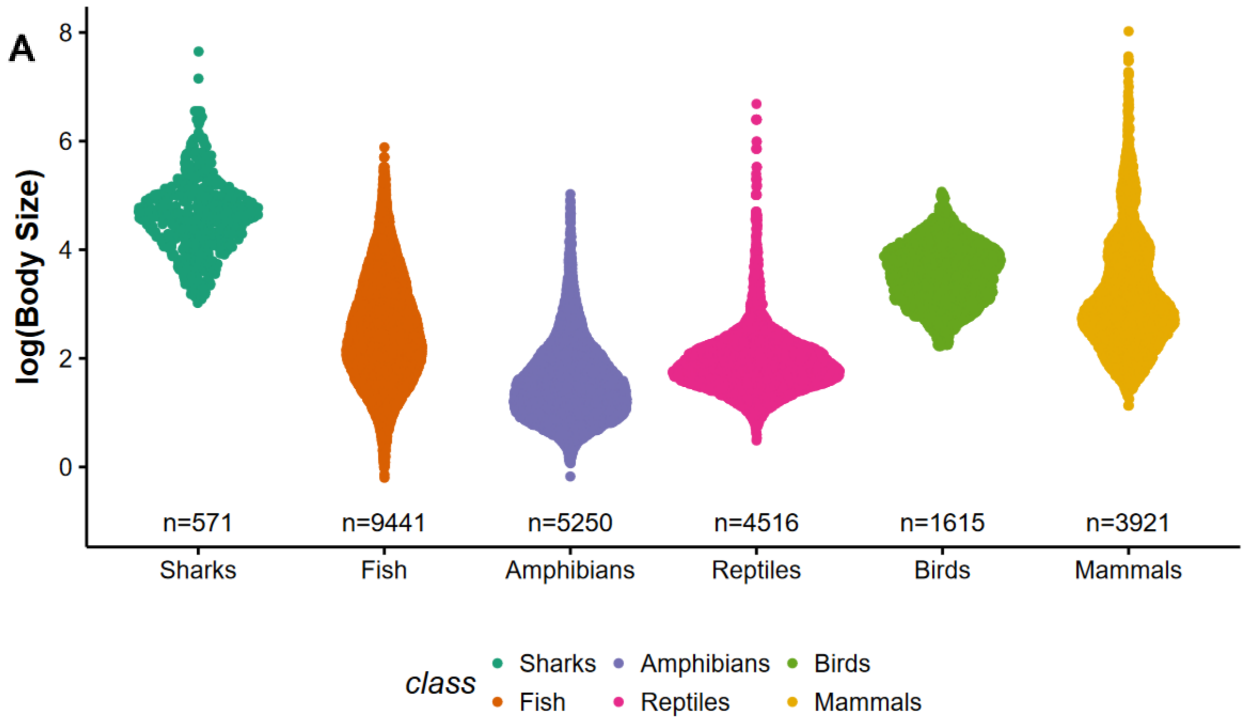


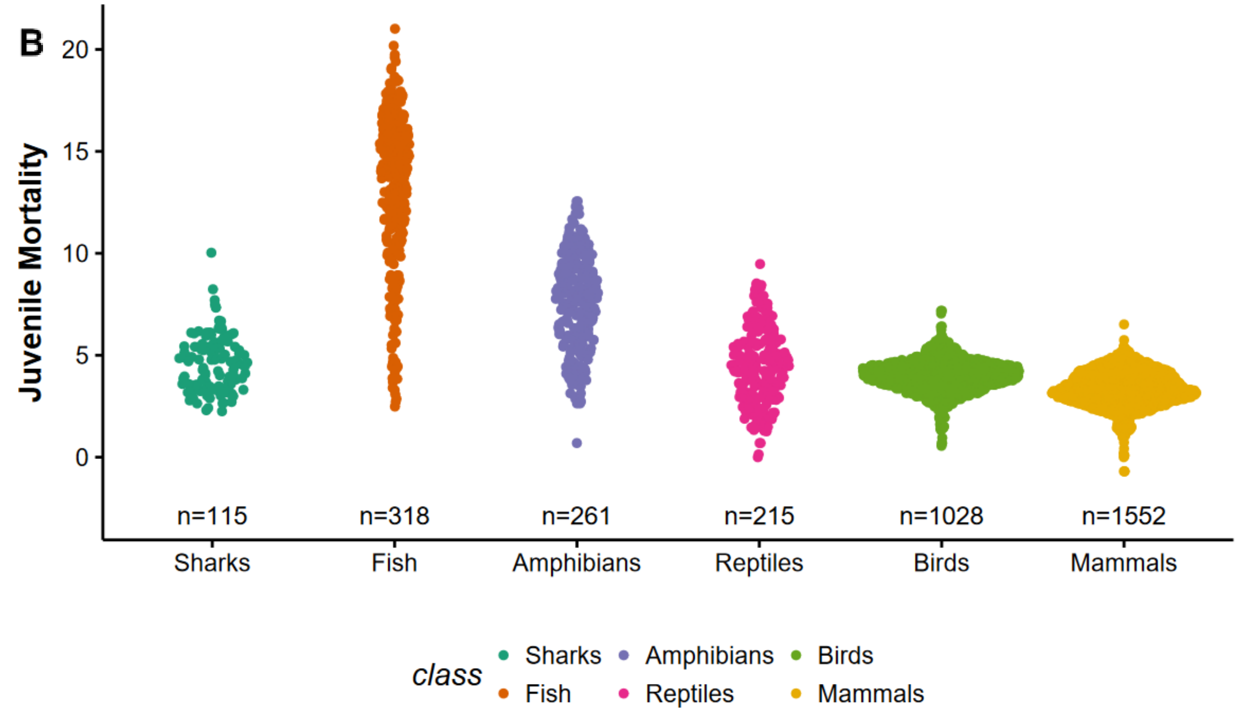

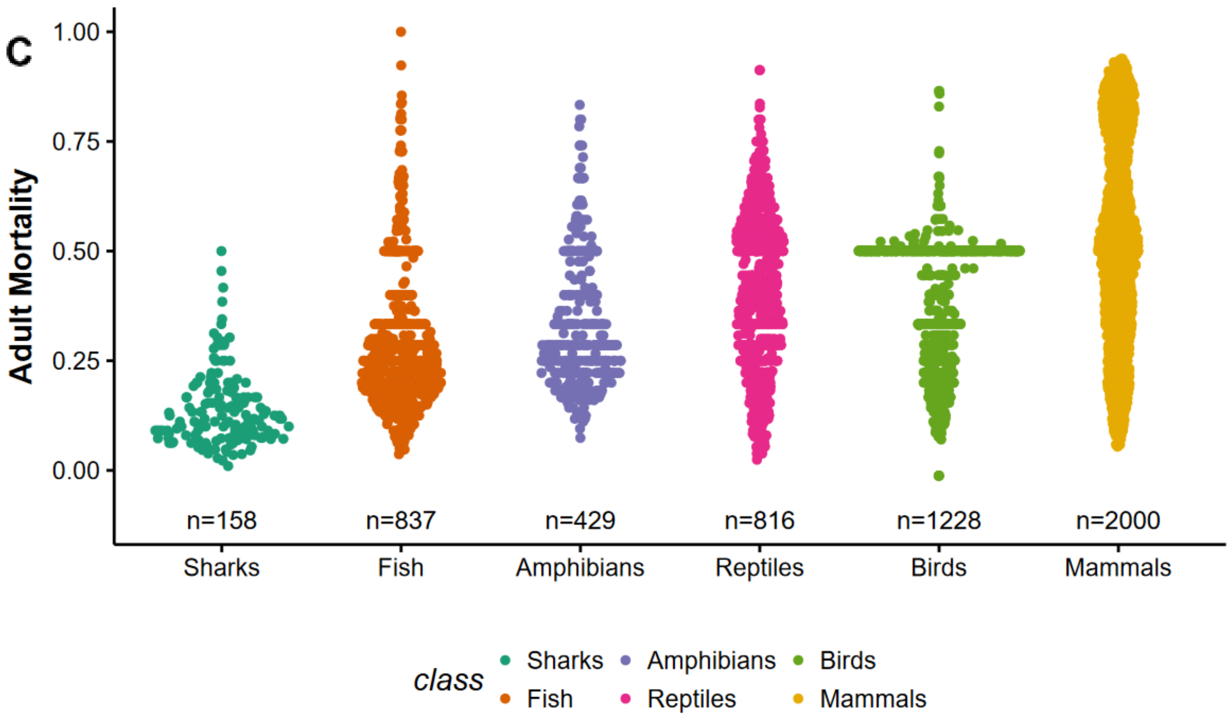


**Supplementary Figure 1: Distribution of vertebrate A) body sizes, B) juvenile mortality, and C) adult mortality.** The number of species in each clade for which trait data is available is provided at the bottom of the panel.

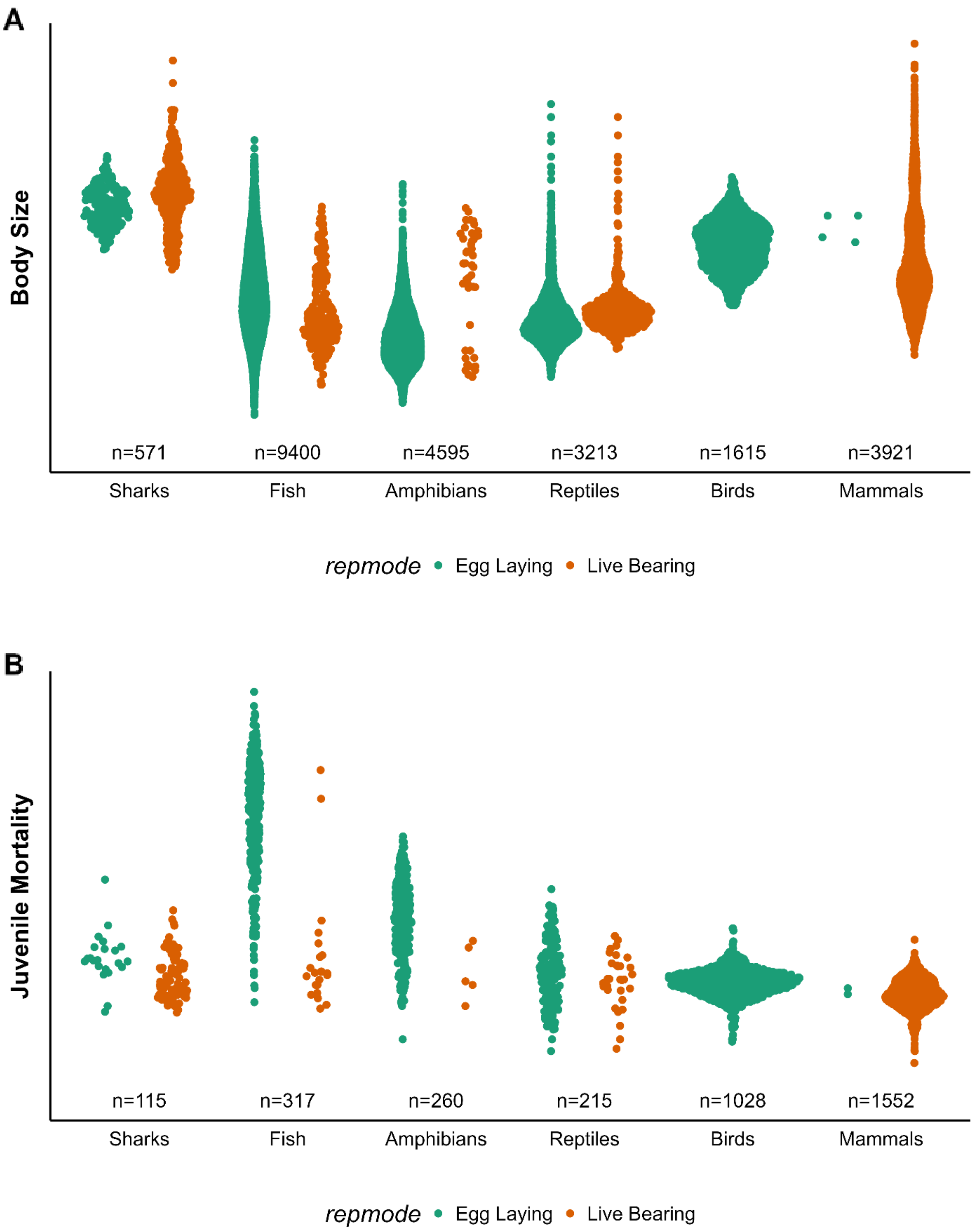
A
Body Size
n=571
n=9400
n=4595
n=3213
n=1615
n=3921
Sharks
Fish
Amphibians
Reptiles
Birds
Mammals
repmode
Egg Laying
Live Bearing
B
Juvenile Mortality
n=115
n=317
n=260
n=215
n=1028
n=1552
Sharks
Fish
Amphibians
Reptiles
Birds
Mammals
repmode
Egg Laying
Live Bearing

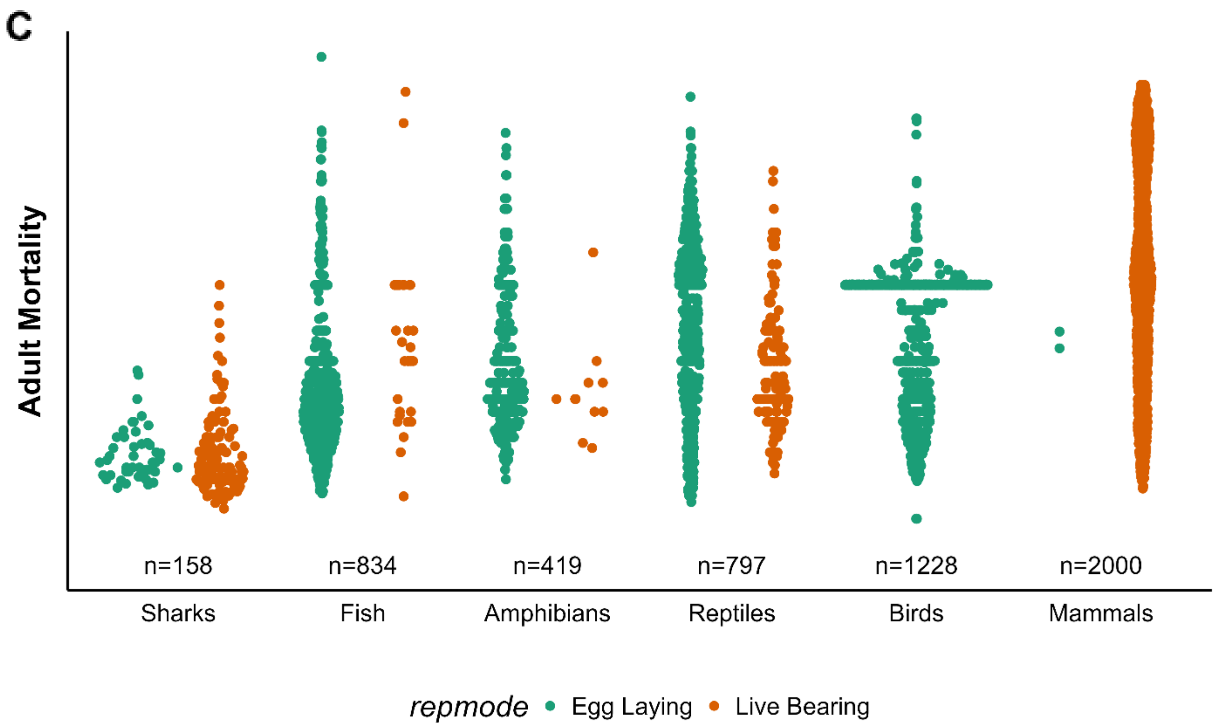


**Supplementary Figure 2: Distribution of egg-laying (turquoise) and live-bearing (orange) vertebrate A) body sizes, B) juvenile mortality, and C) adult mortality.** Points are colored by reproductive mode. The number of species in each clade for which trait data is available is provided at the bottom of the panel.

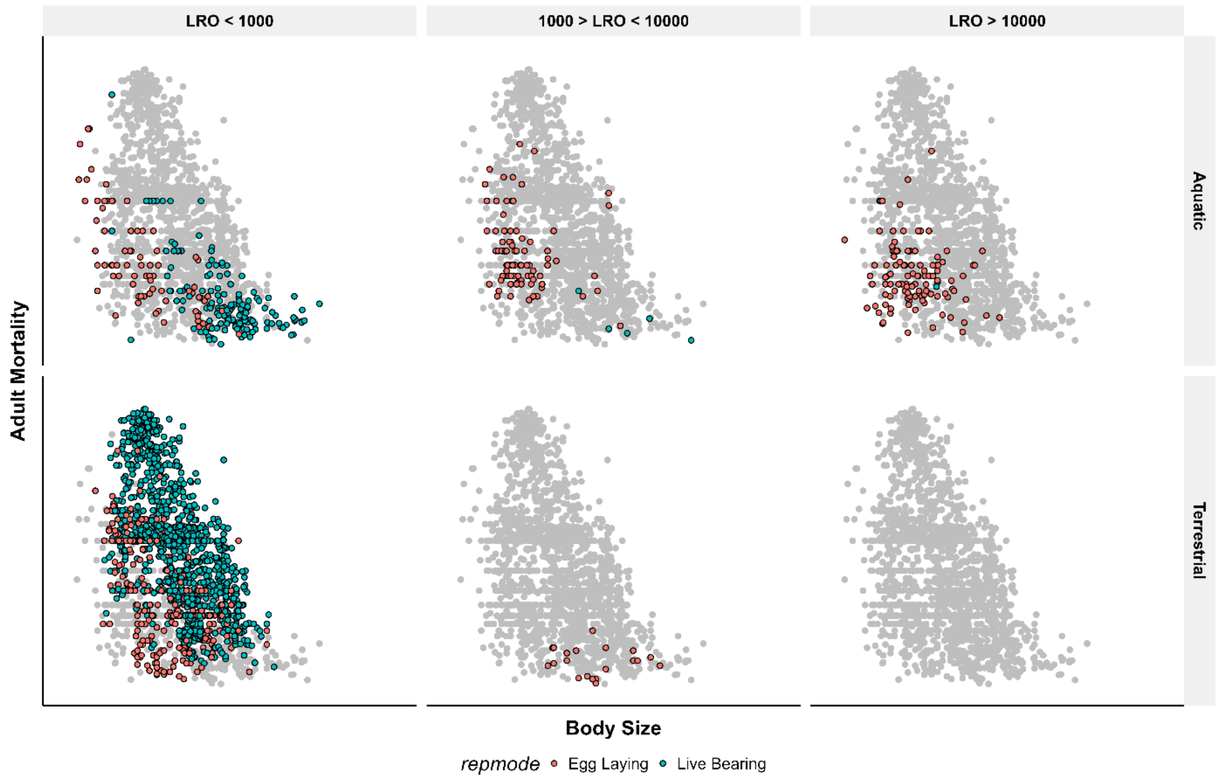


**Supplementary Figure 3. Distribution of aquatic and terrestrial species in relation to body size and adult mortality, for different levels of lifetime reproductive output (LRO).** Adult mortality is estimated as the inverse of age at maturity, and lifetime reproductive output is the raw product of clutch size, longevity minus age at maturity, and the frequency of reproductive bouts. Points are colored by reproductive mode.

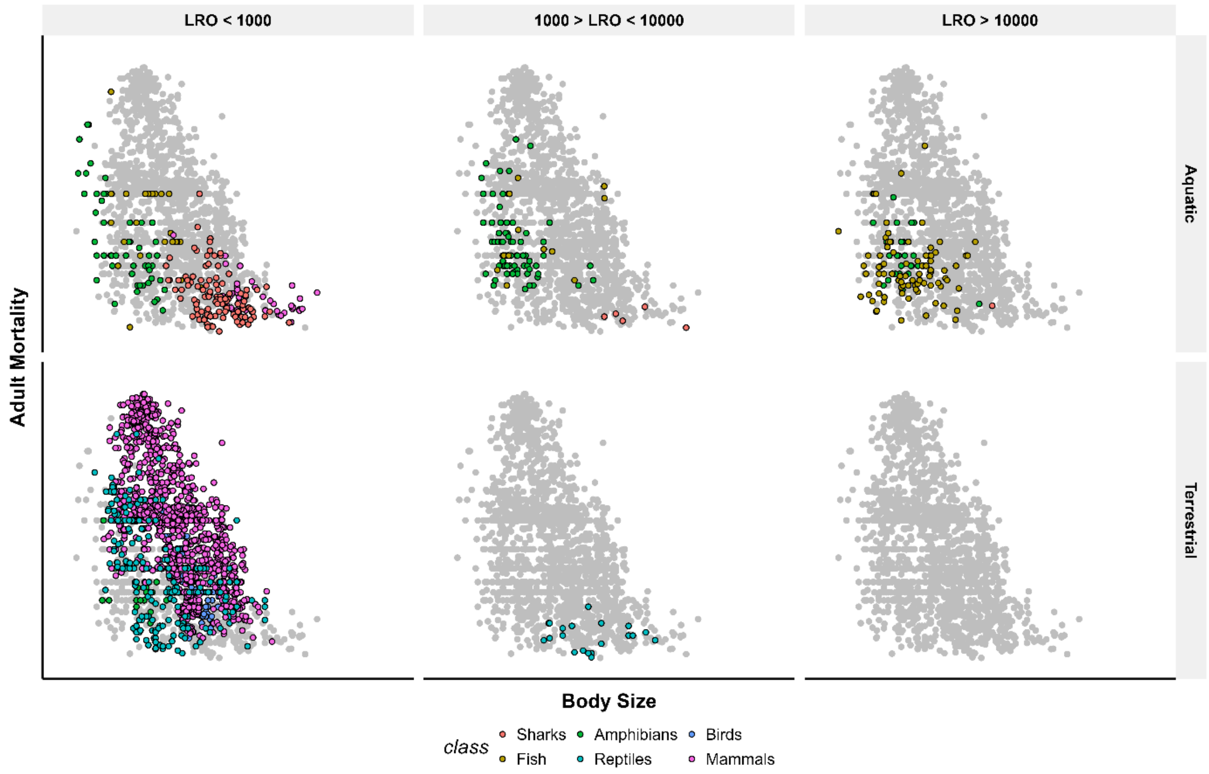


**Supplementary Figure 4. Distribution of aquatic and terrestrial species in relation to body size and adult mortality, for different levels of lifetime reproductive output (LRO).** Adult mortality is estimated as the inverse of age at maturity, and lifetime reproductive output is the raw product of clutch size, longevity minus age at maturity, and the frequency of reproductive bouts. Points are colored by vertebrate class.

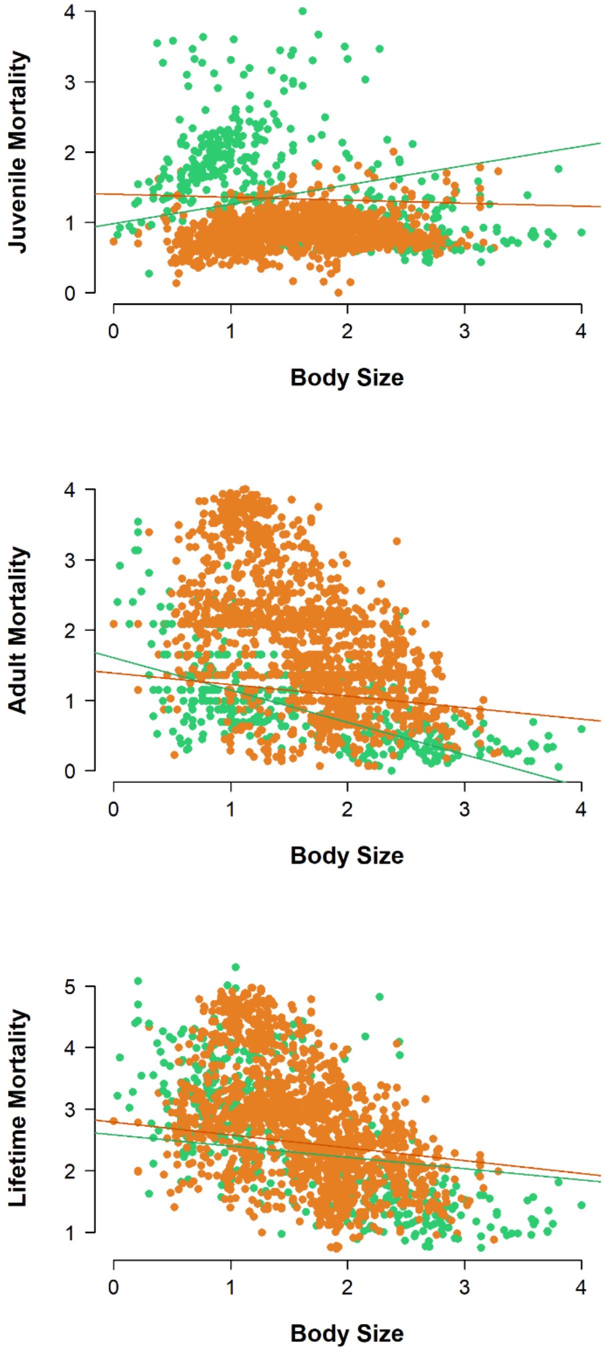


**Supplementary Figure 5: The relationship between body size and a) juvenile mortality, b) adult mortality, and c) lifetime mortality for aquatic (green) and terrestrial (orange) species.** This figure is equivalent to figure 3 in the main text except that predictions from the linear regressions are corrected for phylogeny using the *phylolm* package.

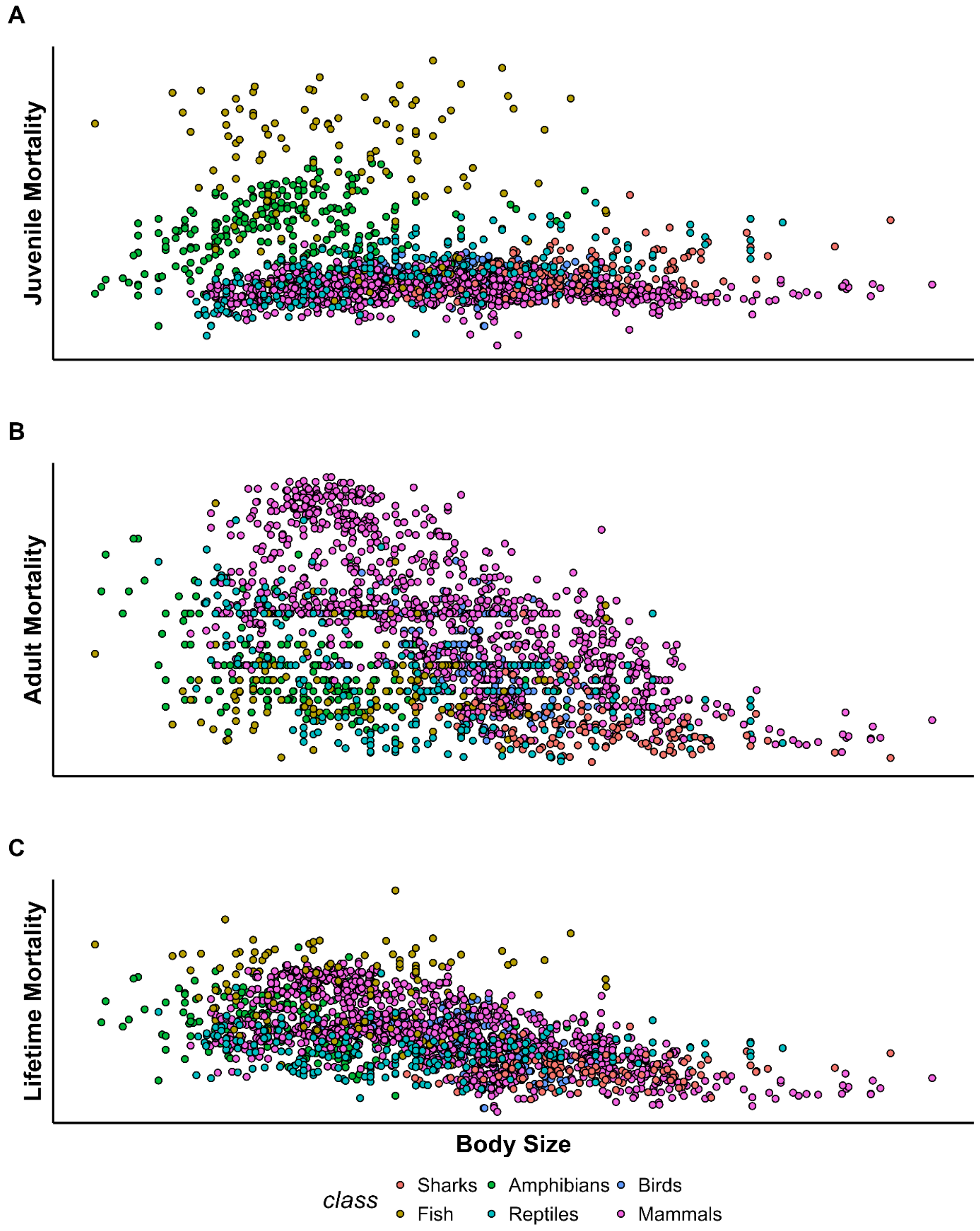


**Supplementary Figure 6: The relationship between body size and A) juvenile mortality, B) adult mortality, and C) lifetime mortality.** Points are color coded by vertebrate class.

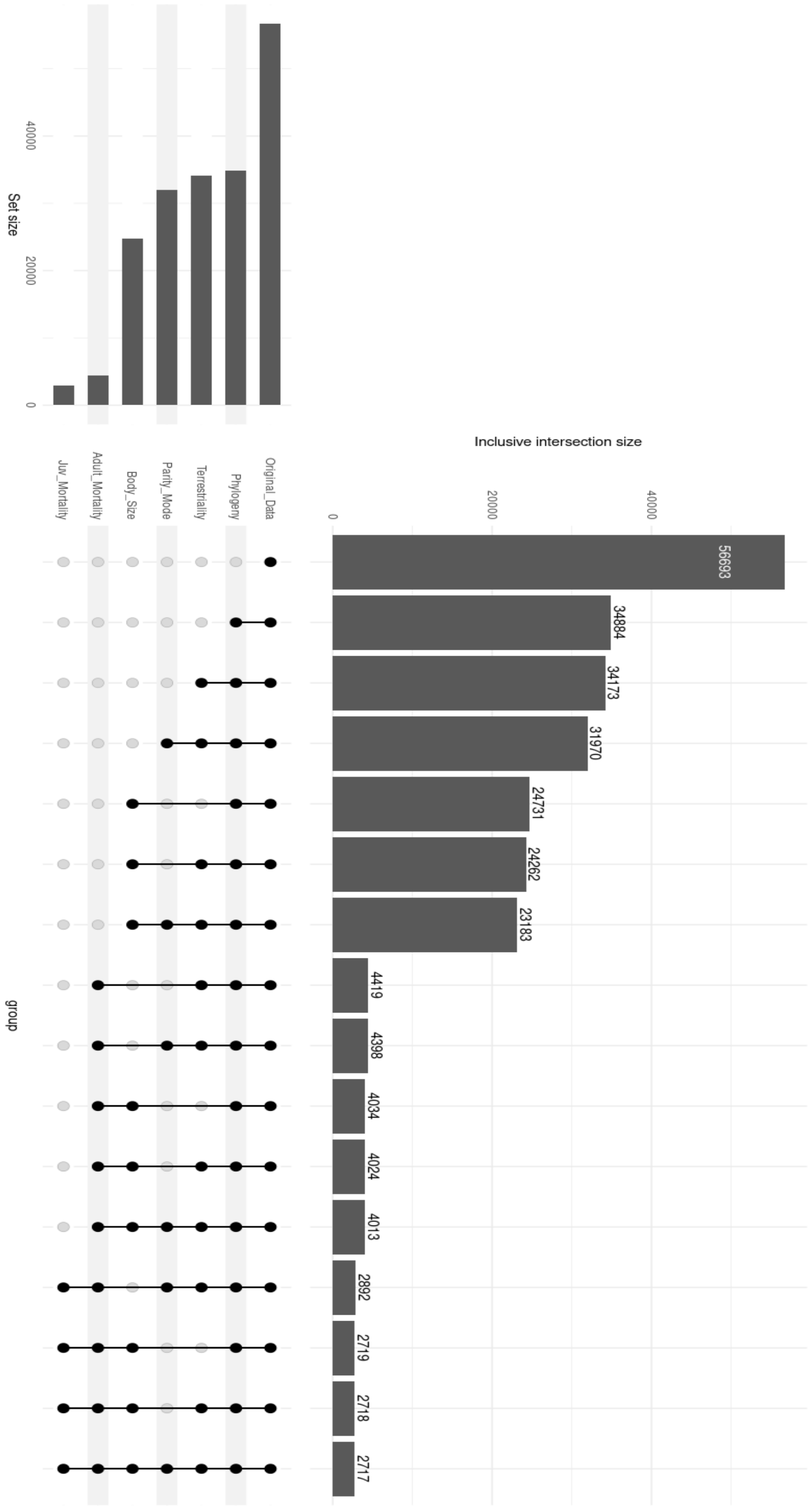


**Supplementary Figure 7. Data coverage across vertebrate clades.** Bars show the proportion of species with published information on terrestriality, parity mode, body size, lifetime reproductive output (juvenile mortality), and age at maturity (adult mortality).

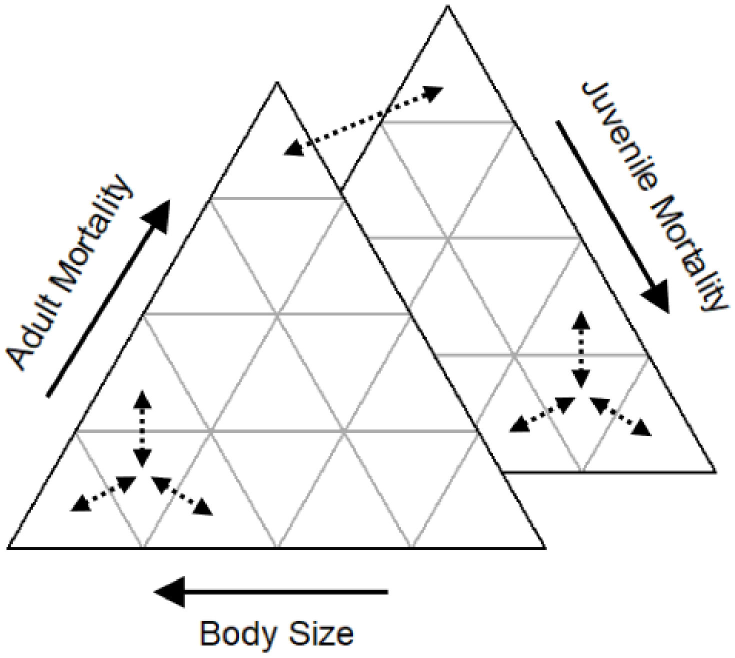


**Supplementary Figure 8. A conceptual diagram of the discrete-space evolutionary model in ternary space.** In this modelling framework, species can evolve to neighbouring patches within the defined life-history space or can transition between states. The two state spaces (triangles) can represent egg-laying and live-bearing reproductive modes or aquatic and terrestrial environments.
